# Mass-spectrometry based proteomics reveals mitochondrial supercomplexome plasticity

**DOI:** 10.1101/860080

**Authors:** Alba Gonzalez-Franquesa, Ben Stocks, Sabina Chubanava, Helle Baltzer Hattel, Roger Moreno-Justicia, Jonas T. Treebak, Juleen R. Zierath, Atul S. Deshmukh

## Abstract

Mitochondrial respiratory complex subunits assemble in supercomplexes. Studies of supercomplexes have typically relied upon antibody-based protein quantification, often limited to the analysis of a single subunit per respiratory complex. To provide a deeper insight into mitochondrial and supercomplex plasticity, we combined Blue Native Polyacrylamide Gel Electrophoresis (BN-PAGE) and mass spectrometry to determine the supercomplexome of skeletal muscle from sedentary and exercise-trained mice. We quantified 422 mitochondrial proteins within ten supercomplex bands, in which we showed the debated presence of complex II and V. Upon exercise-induced mitochondrial biogenesis, non-stoichiometric changes in subunits and incorporation into supercomplexes was apparent. We uncovered the dynamics of supercomplex-related assembly proteins and mtDNA-encoded subunits within supercomplexes, as well as the complexes of ubiquinone biosynthesis enzymes and Lactb, a mitochondrial-localized protein implicated in obesity. Our approach can be applied to broad biological systems. In this instance, comprehensively analyzing respiratory supercomplexes illuminates previously undetectable complexity in mitochondrial plasticity.

**Highlights:**

- Comprehensive quantification of respiratory subunits within supercomplexes
- Complex II and V assemble within supercomplexes
- Mitochondrial-encoded subunits display elevated upregulation upon exercise training
- Exercise increases ubiquinone biosynthesis enzyme complexes

## Introduction

Energy production through oxidative phosphorylation is the primary function of mitochondria. Oxidative phosphorylation is the process by which adenosine triphosphate (ATP) is formed via the transfer of electrons through the electron transport system. The electron transport system is composed of four respiratory complexes (CI to CIV) coupled with ATP-synthase complex (CV). Electrons from the electron carriers NADH and FADH_2_, which are reduced during glycolysis, tricarboxylic acid cycle and beta-oxidation, enter the electron transport system via CI and CII, respectively. Electrons flow through coenzyme Q/ubiquinone, CIII, cytochrome c, and CIV, resulting in pumping of protons into the intermembrane space. The resultant chemiosmotic proton-motive force is used by CV to generate ATP from ADP. Thus, mitochondria play an essential role in cellular metabolism.

The organization of the electron transport system has long been debated, with several models proposed. The *random-collision* model or *fluid* model stated that individual respiratory complexes are free to diffuse within the inner mitochondrial membrane (Hackenbrock et al., 1986). Later, the *solid* model, in which complexes were rigidly superassembled into supercomplexes, was proposed (Schagger and Pfeiffer, 2000). Presently accepted is the *plasticity* model, which takes into account the co-existence of both organisations. In this model, respiratory complexes can dynamically exist in isolation and as supercomplexes depending upon the metabolic requirements of the cell (Acin-Perez et al., 2008). Furthermore, functional respiration in CI-III-IV-containing respirasomes has been confirmed (Acin-Perez et al., 2008; Schagger and Pfeiffer, 2000). Supercomplexes are highly conserved and have been identified in several kingdoms of living organisms (i.e., plants, algae, fungi, protozoa and animals) (reviewed in (Chaban et al., 2014)). The organization of supercomplexes is crucial for individual subunit-complex stability (Acin-Perez et al., 2008), but is also important to reduce ROS production by facilitating electron transport across the complexes (Genova and Lenaz, 2014). Clinically, individuals with genetic mutations in mitochondrial respiratory CI (Ugalde et al., 2004) and CIII (Acin-Perez et al., 2004) present disturbances in supercomplex assembly and stability. Moreover, decreased CI-, CIII- and CIV-containing supercomplexes and mitochondrial respiration have been observed in people with type 2 diabetes (Antoun et al., 2015) and rodent models of aging (Lombardi et al., 2009). CIII and CIV assembly into supercomplexes has also been correlated to substrate utilization and maximal oxygen uptake in humans (Greggio et al., 2017). Hence, respiratory supercomplex formation/dynamics is a physiologically and clinically relevant process.

The formation and plasticity of supercomplexes remain unclear, primarily due to limitations in methodological approaches. Techniques to study respiratory supercomplex formation have typically relied upon Blue Native Polyacrylamide Gel Electrophoresis (BN-PAGE) coupled to antibody-based detection techniques (i.e., oxidative phosphorylation cocktail antibody), often covering only a single subunit per complex. However, this approach provides limited information, relying upon a single subunit to provide representative information regarding complex incorporation into supercomplexes. In the absence of additional targeted analyses, these conventional approaches may also disregard the influence of additional proteins present within supercomplexes, including assembling factors and soluble electron carriers. To date, different studies have combined one- or two-dimensional BN-PAGE with mass spectrometry to study individual mitochondrial respiratory complexes using MALDI-TOF (Farhoud et al., 2005; Sun et al., 2003) or tandem LCMS/MS (Fandino et al., 2005; Wessels et al., 2009). Furthermore, a BN-PAGE LCMS/MS approach was used to cut high-molecular weight sections (not individual bands) to either validate immunoblotting analyses (Greggio et al., 2017) or decipher the composition of individual complexes through agglomerative clustering based on profile-similarity, namely complexome profiling (Guerrero-Castillo et al., 2017; Heide et al., 2012; Van Strien et al., 2019). Thus, while these studies have established a workflow from BN-PAGE to mass spectrometry, this technique has yet to be applied to the analysis of individually targeted supercomplexes.

Mitochondria are highly plastic organelles, rapidly adapting to the metabolic demands of the cell. Exercise training is a potent stimulus to increase mitochondrial respiration within skeletal muscle (Holloszy, 1967) and, therefore, provides an ideal stimulus to study mitochondrial plasticity and supercomplex formation. Using antibody-based detection, supercomplex formation was recently shown to increase within skeletal muscle following exercise training in humans (Greggio et al., 2017). Altered stoichiometry of respiratory complexes within supercomplexes, particularly a redistribution of CI, CIII, and CIV into supercomplexes was identified following exercise training (Greggio et al., 2017). Thus, endurance exercise provides an ideal model to study plasticity in mitochondrial supercomplexes.

In this study we applied an antibody-independent methodology that couples BN-PAGE and LCMS/MS to assess the mitochondrial supercomplexome. Particularly, we utilize exercise training in murine skeletal muscle to study mitochondrial plasticity and supercomplex formation.

## Results and discussion

### Exercise increases mitochondrial respiratory protein subunits in a non-stoichiometric manner

To study mitochondrial plasticity, female C57BL/6JBomTac mice were provided with free access to voluntary wheel running for 25 days or remained sedentary. Immunoblot analysis revealed the expected exercise training-induced increase in hexokinase II and cytochrome c abundance within various skeletal muscle groups (Fig S1A). The greatest exercise training-induced adaptations were apparent in triceps; therefore, this muscle group was selected for subsequent analyses.

Proteomic analysis of skeletal muscle is challenging due to the presence of highly abundant contractile proteins hindering the detection of low abundant proteins (Deshmukh et al., 2015b). We applied a previously established sequential multi-enzyme digestion filter-aided sample preparation (MED-FASP) strategy (with LysC and trypsin) and quantified 3547 proteins within triceps muscle (Schonke et al., 2018; Wisniewski and Mann, 2012) (Fig 1A, Table S1). This strategy allows separation and identification of orthogonal populations of peptides, resulting in increased sequence coverage and depth of the proteome. Principal component analysis (PCA) of quantified proteins revealed clear separation between sedentary and exercise-trained groups (Fig 1B). Amongst the 3547 quantified proteins, 951 were increased and 110 were decreased after exercise training (Fig 1C). The list of significantly increased proteins contains canonical markers of exercise-training such as hexokinase 2 (Hk2), pyruvate dehydrogenase kinase 4 (Pdk4) and lactate dehydrogenase D (Ldhd) (Fig 1C). As expected, mitochondrial protein content was increased after exercise training (Fig 1D), as well as proteins from oxidative phosphorylation, tricarboxylic acid cycle (TCA) and lipid metabolism pathways (Fig 1E). Furthermore, Golgi protein content also increased (Fig 1D). Conversely, proteins from glycolysis and carbohydrate metabolism pathways were downregulated (Fig 1E). This proteomic data confirms the increase in oxidative phosphorylation and shift towards lipid metabolism following endurance exercise training (Constable et al., 1987; Holloszy, 1967; Holloszy and Oscai, 1969; Holloszy et al., 1970; Mole et al., 1971). Proteins involved in substrate transport (Fig S1C), the malate-aspartate shuttle (Fig S1E), the pyruvate dehydrogenase complex (Fig S1H) and the TCA cycle (Fig S1I) all showed general increases. Moreover, proteins with transit peptide (shuttled to mitochondria) (Fig S1J) as well as citrate synthase activity (Fig S1K) were increased upon exercise training. Exercise training also drove an increase in slow-twitch type I and fast-twitch type IIa myosin heavy chain (MHC) isoforms, along with a decrease in the faster fiber type protein MHC IIb (Fig S1L). Thus, via the deepest exercise training-induced skeletal muscle proteome to date, we demonstrate a robust metabolic adaptation to endurance exercise.

**Figure 1.**
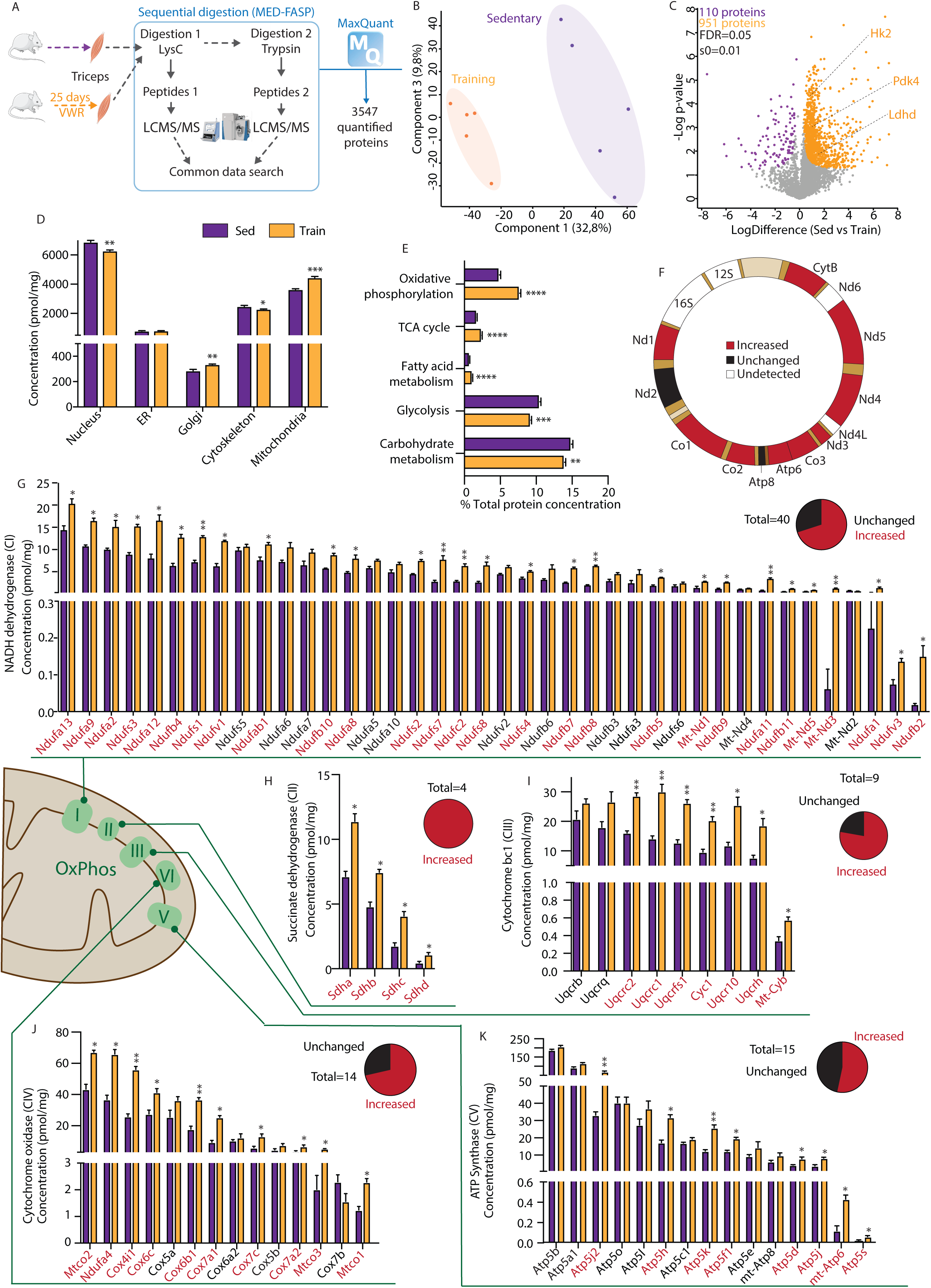
Effects of exercise training on skeletal muscle proteome. A: Experimental design for the study. C57BL\6JBomTac mice were single-housed with or without access to wheel running for 25 days (n=5 per group). Triceps were collected and MED-FASP sample preparation was performed prior to LC-MS/MS analysis. B: Principal component analysis (PCA) segregates the two experimental groups. C: Volcano plot shows all the quantified proteins (110 decreased and 951 increased proteins after exercise). D: Concentration of proteins annotated to different cellular compartments (GOCC). E: Percentage of protein abundance annotated to different molecular functions (GOBP, KEGG for Oxidative phosphorylation). F: Mitochondrial-encoded proteins identified in the proteome, indicated as increased (red), unchanged (black) or undetected (white). G-J: Protein concentration in the triceps proteome for subunits of NADH-dehydrogenase (CI, G), succinate dehydrogenase (CII, H), cytochrome bc1 (CIII, I), cytochrome oxidase (CIV, K), and ATP synthase (CV, J). Complex subunits labeled in red are increased upon exercise training, whilst those in black displayed no significant change. Pie charts in G-J show the proportion of subunits increased/unchanged per complex. Sed: sedentary; Train: exercise trained. All data is represented as mean ± SEM. *p<0.05, **p<0.01, ***p<0.001, ****p<0.0001.

Due to the profound exercise training-induced mitochondrial biogenesis, we studied the mitochondrial part of the proteome in greater detail. Within the total proteome, we detected 11 out of 13 mitochondrial DNA-encoded proteins of which 9 were increased after exercise training (Fig 1F). Indeed, we have excellent coverage across the electron transport system, with 40 subunits in CI, 4 in CII, 9 in CIII, 14 in CIV and 15 in CV quantified within the proteome (Fig 1G-K). Although the majority of proteins increased with exercise training, in most of the complexes, more than one third remained unchanged (Fig 1G-K pie charts: 30% in CI, 22% in CIII, 29% CIV and 47% CV), suggesting altered stoichiometry of respiratory subunits upon exercise training. Although a similar association was observed previously with increased physical activity level (Ubaida-Mohien et al., 2019), the reason for this non-stoichiometric increase in expression has yet to be clarified. Due to the apparent changes in subunit stoichiometry, we questioned the validity of inferring adaptations to whole respiratory complexes from the analysis of a single subunit. Thus, we studied respiratory supercomplex formation using an integrated BN-PAGE mass-spectrometry approach.

### Methodological approach to supercomplexome analysis

The emerging field of protein complexome analyses has paved the way for combining BN-PAGE and LCMS/MS (Guerrero-Castillo et al., 2017; Heide et al., 2012; Van Strien et al., 2019). The predominant approach has consisted of analyzing a large number of continually sliced equal-sized bands (60 slices per sample) along the gel (Guerrero-Castillo et al., 2017; Heide et al., 2012; Van Strien et al., 2019). This allows for the observation of protein distributions across molecular masses. Agglomerative clustering assigns proteins to putative complexes based on similar distribution profiles (Giese et al., 2015; Heide et al., 2012; Van Strien et al., 2019) and thus relies on the assumption that the individual proteins within each supercomplex possesses constant stoichiometries. Furthermore, this requires extensive mass spectrometry measurement time. Given the non-stoichiometric change in respiratory subunits within the triceps proteome we adapted this methodology to excise 10 bands representing known supercomplexes (Fig 2A-B) (Jha et al., 2016) and to allow for comparisons of protein abundance within distinct bands (i.e. supercomplexes). To determine the distribution of proteins across the bands, we appropriated a method for unbiased assignment of proteins to different fractions (traditionally subcellular localizations), namely protein correlation profiling (PCP) (Andersen et al., 2003; Foster et al., 2006; Krahmer et al., 2018). Collectively, this approach allows us to simultaneously assess the abundance and distribution of proteins, specifically within supercomplexes.

**Figure 2.**
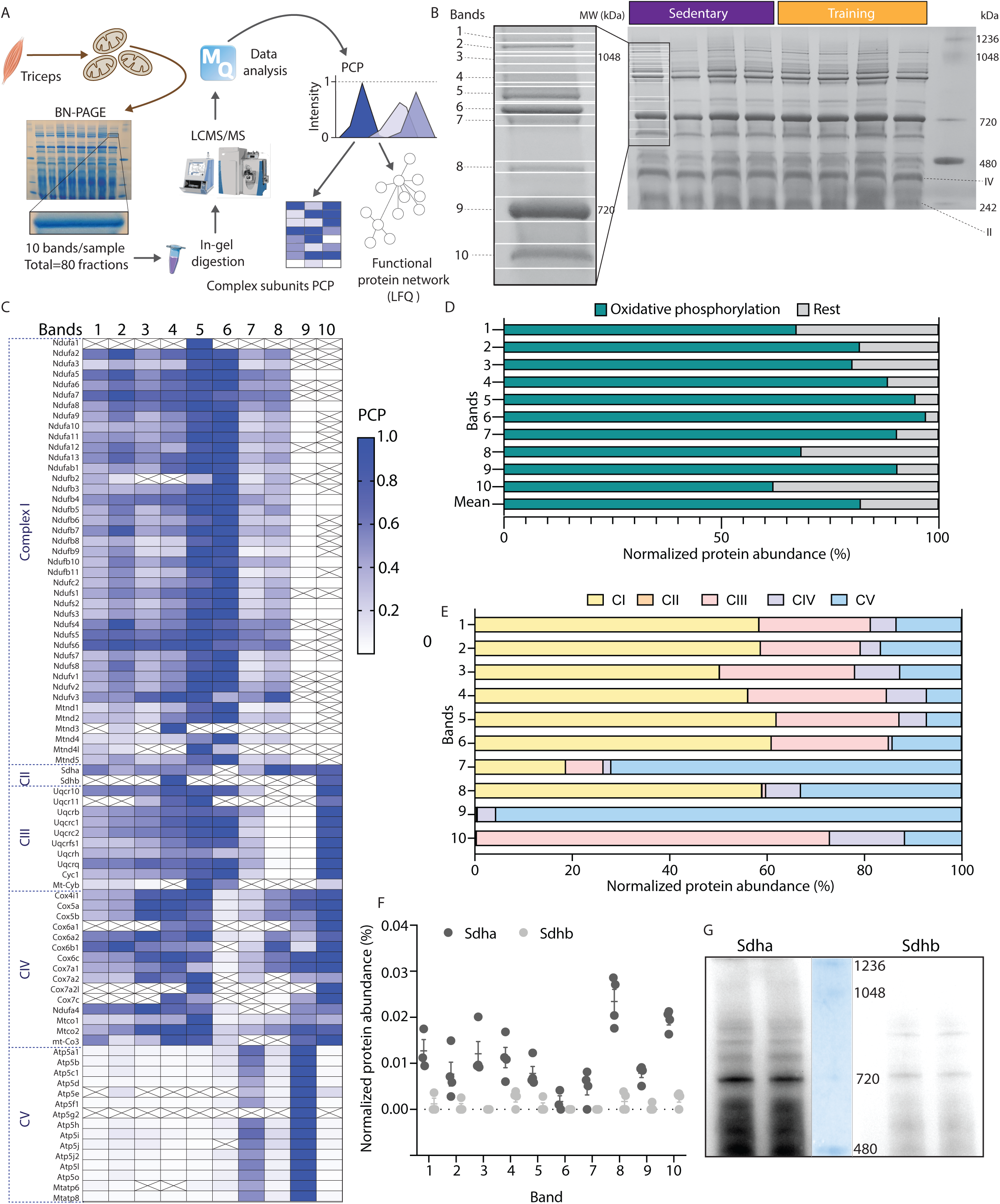
Assessment of mitochondrial supercomplexes by BN-PAGE and LCMSMS. A: Experimental workflow for the study of supercomplex proteomics: mitochondria were isolated from triceps skeletal muscle (n=4 per group) and run in a BN-PAGE gel. Individual bands representing supercomplexes were excised and digested by in-gel digestion prior to LC-MS/MS measurements and data analysis. Protein correlation profile (PCP) of each protein was calculated, and protein-protein interaction networks were generated for some of the bands. B: BN-PAGE image with the different individual bands that were excised per sample. C: PCP heatmap with all the quantified mitochondrial complex subunits within sedentary mice. D: Percentage abundance of oxidative phosphorylation proteins within individual bands (oxidative phosphorylation LFQ intensity within each band is normalized by total LFQ intensity of that band; mean for sedentary group is represented). E: Summed percentage abundance of proteins making up each mitochondrial complex within each band. F: Percentage abundance of Sdha and Sdhb within each band (Sdha and Sdhb LFQ intensity within each band is normalized by total band LFQ intensity of that band; mean ± SEM for sedentary group is represented). H: BN-PAGE immunoblot for Sdha and Sdhb.

### Previously undescribed respiratory subunits within supercomplexes revealed by blue native-PAGE mass-spectrometry

The mitochondrial fraction isolated from triceps muscles was separated on a BN-PAGE gel and bands, representing supercomplexes (Jha et al., 2016), labeled 1-10 were excised after Coomassie blue staining (Fig 2A-B). In-gel digestion (Shevchenko et al., 2006) and LCMS/MS analysis of each band resulted in the quantification of 422 proteins across all bands. We quantified 41 subunits of CI, 2 of CII, 10 of CIII, 15 of CIV and 15 of CV with high confidence (median unique peptides: 6) across 10 of the supercomplex bands (Fig 2C). In support of the methodology, the vast majority of proteins quantified are electron transport system proteins (Fig 2D), validating the excision of respiratory supercomplex bands. As expected (Jha et al., 2016), we observed the presence of CI subunits in the higher molecular weight bands (bands 1-8), while CIV was identified across the majority of supercomplexes (Fig 2C and E). CV showed substantially greater abundance in band 9, making up 95% of the electron transport system protein abundance within this band (Fig 2C and E). These results are in accordance with previous studies, where oligomers of CV subunits were detected in a similar region (Greggio et al., 2017; Jha et al., 2016). We detected subunits of CV within all of the bands analyzed (Fig 2C and E), making up approximately 7-16% of the total electron transport system protein abundance within bands 1-6 (Fig 2E). This agrees with a recent study employing cross-linking mass spectrometry to identify interactions within mitochondrial supercomplexes, which found that CV exists in close spatial proximity, and may physically interact, with known respirasome complexes (Liu et al., 2018).

Strikingly, we quantified the CII subunit succinate dehydrogenase subunit A (Sdha) within the majority of supercomplex bands (Fig 2C and F). The presence of CII within supercomplexes has been debated since CII has not been detected within supercomplexes in all studies (Greggio et al., 2017; Lapuente-Brun et al., 2013; Lenaz and Genova, 2007). However, CII incorporation into superstructures (Kulawiak et al., 2013) and associations with other complexes, namely within supercomplexes, also containing CI, III and IV, have been reported (Acin-Perez et al., 2008; Liu et al., 2018). Furthermore, increased succinate-induced (i.e., CII) respiration has been identified within these supercomplexes (Acin-Perez et al., 2008). In our study, Sdha was quantified in most of the bands (with the exception of band 6; median unique peptides: 8), while Sdhb was only quantified in bands 4 and 10 (median unique peptides: 2; Fig 2C), albeit at a much lower intensity than Sdha (Fig 2F). Accordingly, Sdha has been described to form a higher number of cross-links with other respiratory complexes than Sdhb (Liu et al., 2018). The presence of Sdha, but not Sdhb, within supercomplexes was confirmed by BN-PAGE immunoblotting using subunit-specific antibodies (Fig 2G). This finding could explain some of the discrepancies in the literature regarding CII assembly into supercomplexes. A previous study using an antibody against the lowly expressed Sdhb failed to identify CII within supercomplexes in mouse skeletal muscle (Lapuente-Brun et al., 2013; Lenaz and Genova, 2007). In contrast, CII has been detected within specific supercomplexes, by an anti-Sdha antibody (Acin-Perez et al., 2008). However, another study did not detect Sdha, via immunoblot analysis or mass spectrometry, in supercomplexes from skeletal muscle of elderly (Greggio et al., 2017), suggesting a species-specific difference in supercomplex formation. Nonetheless, functional respiration through CII has only been reported in high-molecular weight murine supercomplexes (Acin-Perez et al., 2008) of approximately similar molecular mass to band 4 identified in the current study, where both Sdha and Sdhb are detected by mass spectrometry. Thus, while Sdha is detectable in most bands, whether these are functional succinate-linked respirasomes remains to be determined. Indeed, Sdhb is necessary for succinate oxidation and electron transfer, while Sdhb knockout reduces oxygen consumption rate (Kitazawa et al., 2017; Rutter et al., 2010). Therefore, the functional role of Sdha in the absence of Sdhb remains unclear.

### Endurance exercise training-induced plasticity in mitochondrial supercomplexes

To study the adaptation of supercomplexes to a stimulus that promotes mitochondrial biogenesis, we assessed the impact of endurance exercise training. The distribution (PCP) of respiratory complexes within each band before and after exercise training is displayed in Fig 3A. The effect of exercise on the distribution of CI, CIII and CV across bands remained broadly similar, while a redistribution of CIV into bands 3 and 4 and a reduction in the proportion of CII within bands 3, 4 and 5 was apparent following exercise training. The mean log_2_-fold-change of the respiratory complexes, as well as complex-related assembly proteins and cytochrome c was calculated (Fig 3B) and absolute changes, rather than distribution across bands, were analyzed. Although, the majority of complexes within each supercomplex show only small changes (typically less than 1 log_2_-fold-change), an increase within CIV and, to a lesser extent CV, within band 7 was notable (Fig 3B). Both of these analyses are consistent with a redistribution of CIV into supercomplexes in skeletal muscle following 4 months of endurance exercise training in elderly sedentary humans (Greggio et al., 2017). Indeed, the proportion of CIV within supercomplexes may be a determinant of energy expenditure during exercise and, thus, central to the energetic adaptation to endurance exercise training (Greggio et al., 2017). However, unlike exercise training in elderly humans (Greggio et al., 2017), a redistribution of CIII was not apparent (Fig 3A), which may reflect differences within species or the training mode/duration.

**Figure 3.**
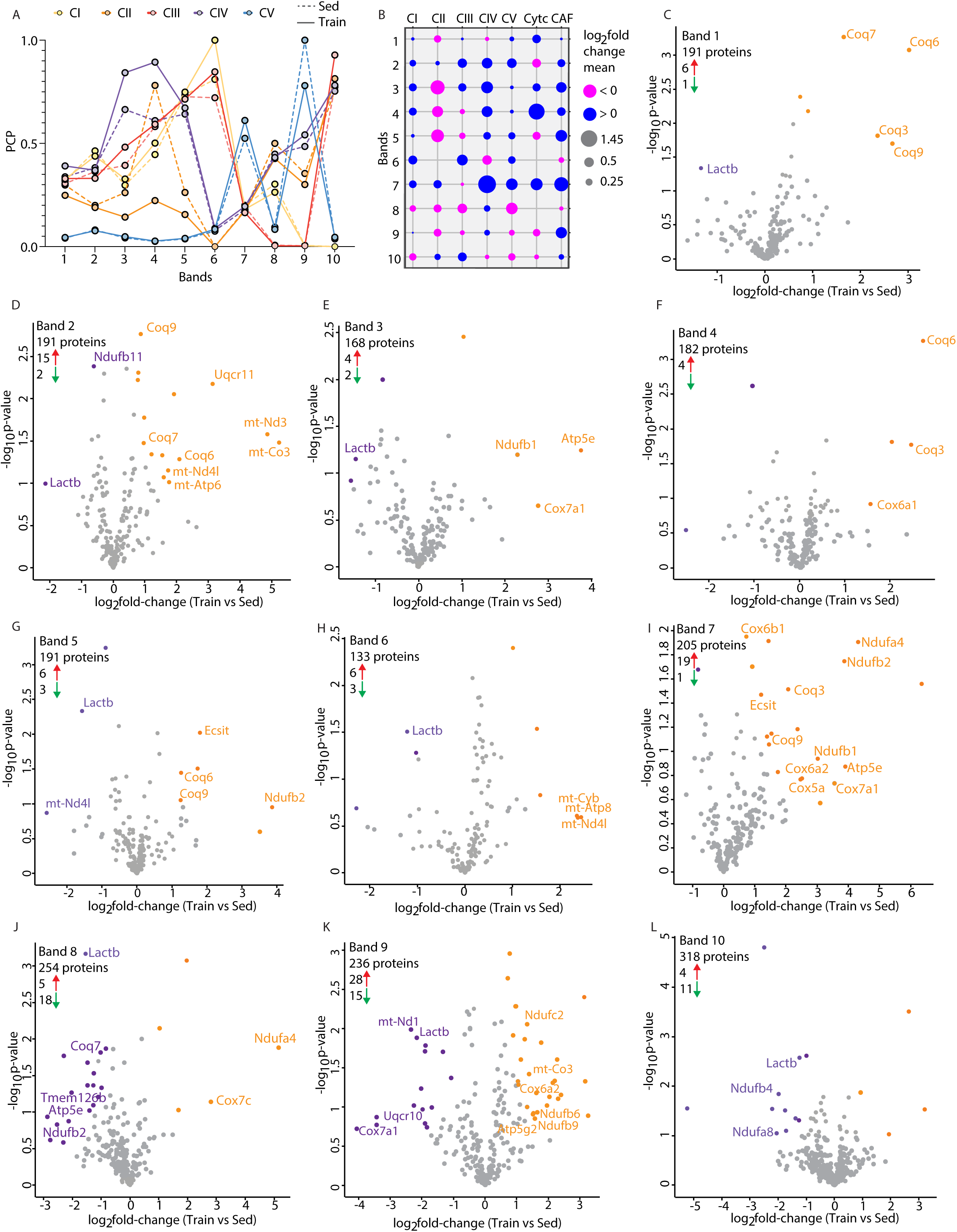
Exercise induces a redistribution of mitochondrial complexes and expression within supercomplexes. A: PCP plot displaying median PCP values of different mitochondrial complexes across the bands (sedentary and trained). B: Bubble chart displaying the mean log_2_-fold-change for each group of oxidative phosphorylation related proteins (mitochondrial complexes, CytC and complex assembly factors – CAF) upon exercise training. C-L: Volcano plots displaying total number of quantified proteins in each band, as well as proteins which were upregulated (red arrow, orange points) and downregulated (green arrow, purple points) in response to exercise in each band (trained vs sedentary). Sed: sedentary; Train: exercise trained.

Analysis of differentially regulated proteins within each band (Figs 3C-L) revealed the adaptation of each supercomplex to exercise training and identified varied changes across bands. For example, band 7 displays the clearest regulation of respiratory subunits following exercise training. Mitochondrial respiratory complex proteins from CI (Ndufa2, Ndufa4, Ndufb1 and Ndufb2), CIV (Cox5a, Cox6a2, Cox6b1 and Cox7a1) and CV (Atp5e) were increased upon exercise training (Fig 3I). Furthermore, the complex assembly factor Evolutionary Conserved Signaling Intermediate in Toll Pathways (Ecsit) was also increased within band 7 (Fig 3I) and band 5 (Fig 3G). Analyses of the volcano plots also identify non-electron transport system proteins that are regulated by exercise training and form complexes within a similar mass range. The Coenzyme Q biosynthetic family (COQs) showed consistent increases across the majority of the bands (bands 1, 2, 4, 5 and 7). Additionally, lactamase beta (Lactb), a mitochondrial localized protein implicated in obesity (Chen et al., 2008) was consistently reduced (bands 1, 2, 3, 5, 6, 8, 9 and 10) following exercise training (Figs 3C-L).

### Potential role of respiratory complex assembly factors in supercomplex assembly

Mitochondrial complex assembly factors play a critical role in the construction of functionally active respiratory complexes, which can organize themselves into supercomplexes (Ghezzi and Zeviani, 2012). We investigated the presence of mitochondrial complex assembly factors (as annotated in HUGO, genenames.org) and quantified 14 within supercomplex bands (Fig 4A). This is in comparison to 35 complex assembly factors quantified within the total triceps proteome (12 for CI, 1 for CII, 5 for CIII, 11 for CIV, 3 for CV; Fig 4C), indicating that many assembly factors may complete their role prior to the formation of supercomplexes. Of note, 10 of the 14 complex assembly factors quantified within supercomplexes are CI assembly factors, reflecting the complicated process of assembling CI from 44 subunits. In general, PCP analyses revealed higher expression of assembly factors within low-molecular mass bands (Fig 4A). This agrees with a previous *in vitro* study whereby almost complete dissociation of CI assembly factors in high-molecular mass supercomplexes was reported (Guerrero-Castillo et al., 2017). The predominant presence of assembly factors within bands 7-10 may indicate these as precursors to higher-molecular mass supercomplexes. The temporal nature of complex assembly and incorporation into supercomplexes has been debated (Acin-Perez et al., 2008; Guerrero-Castillo et al., 2017; Moreno-Lastres et al., 2012). Using an antibody-based technique *in vitro*, an incomplete sub-assembly of CI forms an approximately 830-kDa complex with partially-assembled CIII and CIV before full assembly of each complex occurs within the formative supercomplex (Moreno-Lastres et al., 2012). However, in a similar model, albeit utilizing mass-spectrometry-based complexome analysis, evidence was provided to show that supercomplex assembly occurs only after full assembly of the individual complexes (Guerrero-Castillo et al., 2017). In our *in vivo* analysis of mature skeletal muscle, we provide evidence that likely supports the full assembly of individual complexes prior to assembly of supercomplexes. Indeed, we found almost all subunits of CI could be detected in each of bands 1-8, including Ndufa6, Ndufa7, Ndufa11 and Ndufab1, which are reported to assemble in the final steps of CI formation (Guerrero-Castillo et al., 2017) (Fig 2C). There was also no evidence of partially assembled CI complexing with CIII or CIV, even in the approximately 830 kDa band 8. Nonetheless, the presence of CI, CII and CIV, with very little CIII detected within band 8 may indicate an intermediate supercomplex, which precedes the formation of the CI, (II), III and IV respirasome. Band 8 showed a general trend of downregulated electron transport system proteins following endurance exercise training (Figs 3B and J; including CI proteins: Ndufa2, Ndufb2, Ndufb11, Ndufs2; CIV protein: Cox7a1; and CI assembly protein: Tmem126b), potentially suggesting a shift towards mature respirasomes during a period of elevated energetic demand.

**Figure 4.**
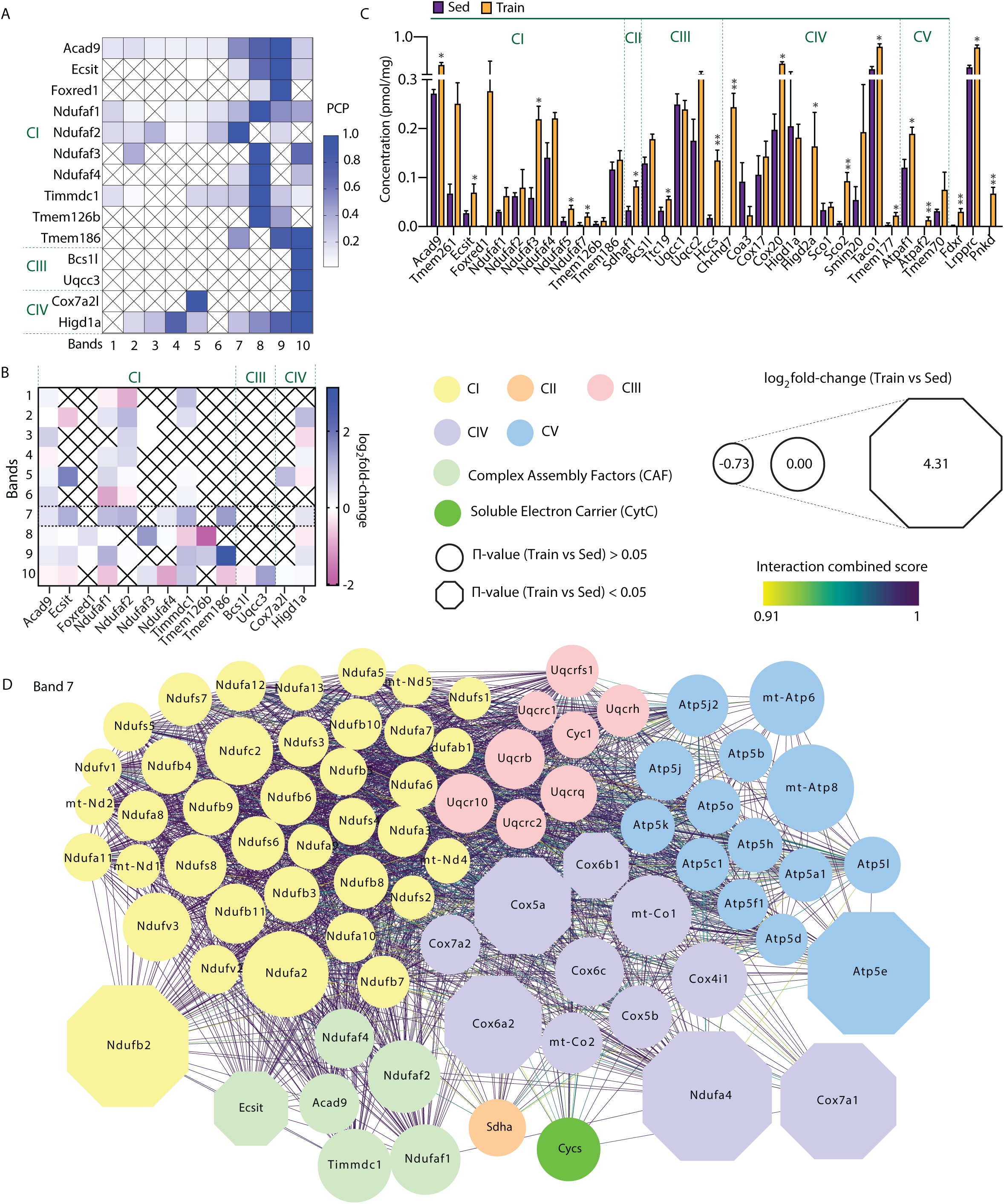
Complex assembly factors in supercomplex regulation. A: PCP heatmap including all the quantified mitochondrial complex subunits within sedentary mice. B: Heatmap representing log_2_-fold-change (trained vs sedentary) for complex assembly factors across all bands. C: Protein concentration in the triceps proteome for complex assembly factors. Data is represented as mean ± SEM. *p<0.05, **p<0.01. C: Protein-protein interaction network (STRING confidence score 0.7) generated from oxidative phosphorylation proteins, CytC and complex assembly proteins in band 7. Edge color indicates the combined score of the interaction, while node size indicates log_2_-fold-change with hexagonal nodes indicating a significantly regulated protein (π value < 0.05; (Xiao et al., 2014)). Sed: sedentary; Train: exercise trained.

The effect of exercise training upon the 14 complex assembly factors within the supercomplexes was determined (Fig 4B). Within band 7, we identified the upregulation of 8 assembly factors following endurance exercise training, including the complex I assembly factor Ecsit (Fig 4B). In addition, 21 assembly factors were upregulated in the total triceps proteome upon exercise training (Fig 4C). This is in agreement with a study that showed assembly factor proteins Ndufaf3, Ndufaf4, Uqcc2, Cox20 and Atpaf1 were associated with higher physical activity in human skeletal muscle (Ubaida-Mohien et al., 2019). Conversely, we did not observe changes in Sco1 and Uqqc1 upon exercise training, which may reflect species-specific or methodological (i.e., observational versus interventional) differences.

Ecsit, when localized in the mitochondria, is involved in the assembly and stability of CI (Vogel et al., 2007). To understand the increase in Ecsit incorporation into some supercomplex bands we studied band 7 in further detail. A protein-protein interaction network (STRING confidence score > 0.7) was generated for band 7 displaying quantified mitochondrial complex proteins, complex assembly factors, and soluble electron carrier related proteins (Fig 4D; Table S4). Increased Ecsit protein abundance was concomitant with upregulated CI, CIV and CV proteins (octagon-shaped). This network revealed 45 high-confidence protein-protein interactions between Ecsit and other proteins, including proteins 38 subunits of CI and numerous CI assembly factors as well as one CIII protein and one CV protein (Fig 4D; Table S3). Ecsit is recruited to mitochondria partly due to a N-terminal mitochondria-targeting sequence (Vogel et al., 2007), where it associates within the mitochondrial CI assembly complex (MCIA) (including Acad 9, Ndufaf1, Timmdc1 and Tmem126b (Guarani et al., 2014; Heide et al., 2012) and with other CI assembly proteins (Ndufaf2 and Ndufaf4). Knockdown of Ecsit results in decreased Ndufaf1 and CI protein abundance, which ultimately leads to mitochondrial dysfunction (Vogel et al., 2007). Ecsit knockdown cells also show an increase in superoxide (Koopman et al., 2005) and cytosolic oxidant levels (Koopman et al., 2006). Collectively, increased Ecsit protein within remodeling supercomplexes (e.g., band 7) may indicate a role for Ecsit in supercomplex assembly over and above the assembly of CI.

### Mitochondrial-encoded subunits display elevated upregulation within supercomplexes following exercise training

Mammalian mtDNA encodes for 13 complex subunits: 7 subunits for CI, 1 for CIII, 3 for CIV and 2 for CV. We identified 11 mitochondrial-encoded respiratory subunits across the different bands, of which a number were upregulated in several bands after exercise training (Fig3 C-L; Fig 5A). On average, mitochondrial-encoded subunits display greater upregulation within supercomplexes than nuclear-encoded subunits for the same complex (Fig 5B) in response to exercise training, indicative of altered stoichiometry of the subunits that form supercomplexes. Band 2 shows upregulation of mitochondria-encoded subunits within CI, CIII, CIV and CV (Fig 5A), in excess of their nuclear-encoded counterparts (Fig 5B). In order to visualize the exercise training-induced regulation of mitochondrial- and nuclear-encoded proteins within supercomplexes, we generated a protein-protein interaction network within band 2 (STRING confidence score > 0.7) (Fig 5D; Table S5). Within this band, particularly for mt-Nd3 and mt-Co3, but also mt-Nd4l and mt-Atp6, exercise training clearly induces a greater upregulation of mitochondrial-encoded than most nuclear-encoded subunits (Fig 5D). Moreover, 11 mtDNA subunits were identified in the whole triceps proteome, with a general upregulated trend upon exercise training (Fig 5C), corroborating previous evidence (Menshikova et al., 2006; Puntschart et al., 1995; Sae-Tan et al., 2014; Yokokawa et al., 2018). These findings suggest that exercise training increases mitochondrial-encoded subunit transcription, consistent with increased mtDNA and mitochondrial genes after different types of exercise (Robinson et al., 2017), or increased mitochondrial-encoded gene translation. The latter is supported by upregulated mitochondrial ribosomal proteins upon exercise training in the whole triceps proteome (Fig 5E). Taco1 (Translational activator of cytochrome c oxidase 1), a protein annotated as an assembly factor, but also a translational activator binding mitochondrial mt-Co1 mRNA and regulating mitochondrial proteins expression (Richman et al., 2016), was increased in the whole triceps proteome (Fig 4B), along with mt-Co1 (Fig 5B). This corroborates a previous observation of voluntary wheel running-induced increase in Taco1 (Lee et al., 2016). Moreover, the increase in mitochondrial mRNA translational machinery coincides with the exercise-induced increase in mitochondrial biogenesis (Yokokawa et al., 2018). Taco1 mutations have been associated with CIV deficiency and mitochondrial dysfunction in human fibroblast primary cells (Weraarpachai et al., 2009) and rodents (Richman et al., 2016). Collectively, our study provides evidence that exercise-training upregulates mitochondrial-encoded subunits within supercomplexes, which is coincidental with an increase in the mitochondrial translational machinery.

**Figure 5.**
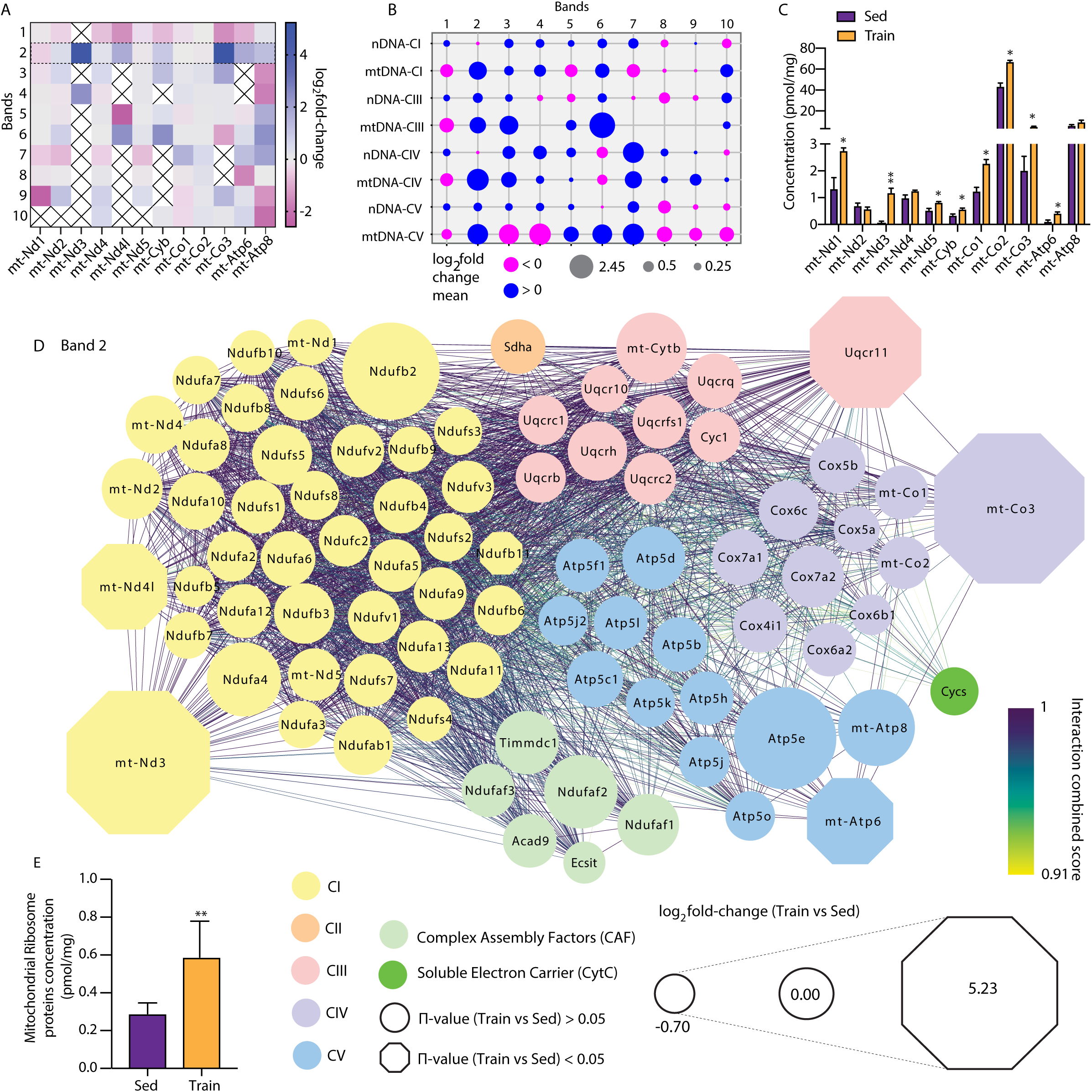
Mitochondrial-encoded oxidative phosphorylation proteins are upregulated in supercomplexes following exercise training. A: Heatmap displaying log_2_-fold-change (trained vs sedentary) for mitochondrial-encoded oxidative phosphorylation proteins across the different bands. B: Bubble chart displaying the log_2_-fold-change for mitochondrial- and nuclear-encoded proteins for each complex within supercomplexes. C: Protein concentration of mitochondrial-encoded oxidative phosphorylation proteins in the triceps proteome. Data is represented as mean ± SEM. *p<0.05, **p<0.01. D: Protein-protein interaction network (STRING confidence score 0.7) generated from oxidative phosphorylation proteins, CytC and complex assembly proteins in band 2. E: Total protein concentration in the triceps proteome for proteins of mitochondrial ribosomal proteins. Sed: sedentary; Train: exercise trained.

### Exercise-training increases the ubiquinone biosynthetic family of COQs

The presence of soluble electron carriers in the electron transport system was proposed decades ago (Green and Tzagoloff, 1966). Ubiquinone transfers electrons from CI or CII to CIII, while CytC transfers electrons from CIII to CIV. Electron carriers have previously been identified in respiratory supercomplexes (Acin-Perez et al., 2008; Althoff et al., 2011). In the current study, we detected CytC within each supercomplex band,with the exception of bands 6 and 8 (Table S2).

Ubiquinone biosynthesis enzymes (COQs) form complexes of 700-1300 kDa (Floyd et al., 2016; Marbois et al., 2009) and, as such, have co-migrated into many of the bands excised for supercomplex analysis. COQs showed substantial increases in abundance across many of these bands (Fig 3C-L & Fig 6A). COQ5, COQ7 and COQ9 also showed a significantly increased concentration in the total triceps proteome, while a similar tendency was apparent for COQ6 (Fig 6B). Thus, COQ enzymes are upregulated concomitant with the increased metabolic demand in response to endurance exercise training. The upregulation of the COQ complex has been demonstrated during galactose-induced mitochondrial biogenesis in HepG2 cells *in vitro* (Floyd et al., 2016), while downregulation of COQ enzymes is apparent in rodent models of mitochondrial dysfunction (Kuhl et al., 2017). However, to our knowledge, this is the first demonstration of *in vivo* COQ protein complex regulation following a mitochondrial biogenic stimulus in mammals. The increase in COQs in response to exercise training may explain the known increase in ubiquinone content in oxidative skeletal muscle following exercise training (Gohil et al., 1987). An increase in COQs is also apparent during mitochondrial biogenesis in liver, as well as white and brown adipose tissue (Aithal et al., 1968; Bentinger et al., 2003; Gohil et al., 1987; Quiles et al., 1994). Upregulation of ubiquinone has therapeutic effects, with exogenous ubiquinone administration increases exercise performance (Alf et al., 2013; Cooke et al., 2008; Ylikoski et al., 1997) and reverses symptoms of many pathophysiological conditions in humans and animals (Di Giovanni et al., 2001; Garrido-Maraver et al., 2014; Quinzii and Hirano, 2010; Rotig et al., 2000; Xu et al., 2010). Future studies should aim to determine whether upregulated COQ enzyme complexes upregulated ubiquinone within specific pools (e.g., free vs supercomplex bound pools).

**Figure 6.**
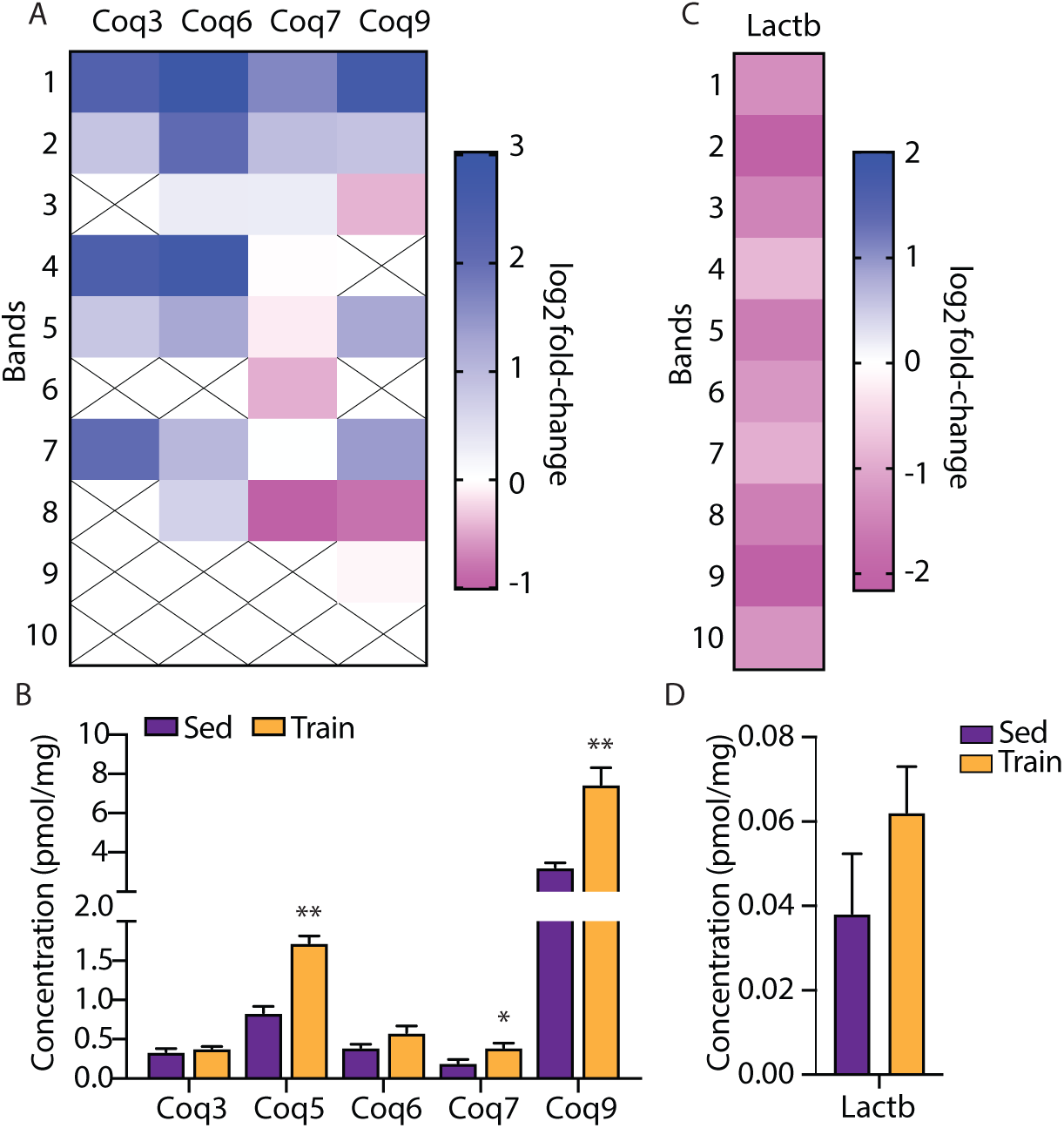
Effect of exercise on non-oxidative phosphorylation protein complexes. A: Heatmap displaying log_2_-fold-change (trained vs sedentary) for ubiquinone biosynthetic family of COQ proteins across the different bands. B: Protein concentration of COQ proteins in the triceps proteome. C: Heatmap displaying log_2_-fold-change (trained vs sedentary) for Lactb protein across the different bands. D: Protein concentration of Lactb in the triceps proteome. Data is represented as mean ± SEM. *p<0.05, **p<0.01. Sed: sedentary; Train: exercise trained.

### Obesogenic Lactb polymers decrease in mitochondria of exercise-trained mice

Our analyses also identified downregulated proteins in all supercomplex bands (Figs 3C-L, Table S2). For instance, Lactb was downregulated within the majority of the bands (Figs 3C-L and Fig 6C). The Lactb gene is positively correlated with obesity (Chen et al., 2008; Yang et al., 2009) and type 2 diabetes (Lau et al., 2017), while Lactb overexpression in transgenic mice increases fat mass (Chen et al., 2008). Lactb is localized in the mitochondrial intermembrane space and polymerizes to form filaments with a molecular mass of >600 kDa (Polianskyte et al., 2009). Therefore, a downregulation of Lactb polymers may confer metabolic advantages following endurance exercise training. Lactb overexpression reduces the phospholipids lipophosphatidylethanolamine and phosphatidylethanolamine in mitochondria of breast cancer cells, but not in non-transformed cells, concomitant with suppression of tumorigenesis (Keckesova et al., 2017). The metabolic function of Lactb in non-tumorigenic cells is unclear. Despite a downregulation of Lactb in high-molecular mass complexes (Fig 3C-L and Fig 6C), Lactb expression in the total proteome shows a non-significant trend to increase (Fig 6D). Thus, polymerization, and not the total content of Lactb, appears to be influenced by endurance exercise training.

## Conclusion

Limitations in methodologies have constrained the comprehensive investigation of mitochondrial supercomplex plasticity. Here, we described a BN-PAGE mass spectrometry approach to study the supercomplexome, which has broad applicability to investigate protein complexes in various systems. We applied this methodology to identify the composition and dynamics of respiratory supercomplexes within skeletal muscle. We quantified 41 subunits of CI, 2 of CII, 10 of CIII, 15 of CIV and 15 of CV across 10 supercomplex bands, identifying the debated CII and CV as components of respiratory supercomplexes. Furthermore, mitochondrial respiratory complex assembly proteins were identified in low molecular-mass supercomplexes, suggesting a role in early supercomplex assembly. We also uncovered the dynamics of the exercise training-induced supercomplexome and identified the mitochondrial-encoded proteins within these supercomplexes. Finally, we also identified additional high-molecular mass complexes, and described the regulation of the ubiquinone biosynthesis enzyme complex and Lactb polymers in response to exercise training. Collectively, this not only highlights the power of utilizing a proteomic approach to study complexes and supercomplexes, but also provides biological insight into the plasticity of mitochondria during endurance exercise-induced mitochondrial biogenesis.

## Acknowledgments

This work is supported by an unconditional donation from the Novo Nordisk Foundation (NNF) to NNF Center for Basic Metabolic Research (http://www.cbmr.ku.dk) (Grant number NNF18CC0034900) and NNF Center for Protein Research (https://www.cpr.ku.dk/) (Grant number NNF14CC001). This work was partially supported by an EFSD/Lilly grant to AGF, Swedish Research Council (2015-00165) and Novo Nordisk Foundation Challenge grants (NNF14OC0011493) to JRZ, and Novo Nordisk Foundation Excellence Project Award (NNF14OC0009315) and the Danish Council for Independent Research (DFF 4004-00235) grants to JTT. We would like to acknowledge Matthias Mann, Erik Richter, Jørgen F.P. Wojtaszewski, Alberto Santos, Ana Rita Colaço, Rebeca Soria and the MS platform at the NNF Center for Protein Research.

## Author’s contributions

ASD and JRZ supervised this work. ASD and AGF: hypothesis generation, conceptual design, data analysis and manuscript preparation. ASD, AGF and BS: Wrote manuscript. ASD, HBH, SC, JTT, AGF, RMJ and BS: performed experiments. JRZ edited the manuscript. All authors reviewed the manuscript. ASD is the guarantor of this work and, as such, had full access to all the data in the study and takes responsibility for the integrity of the data and the accuracy of the data analysis.

## Context and Significance

Mitochondria are the powerhouses of the cell, producing the majority of energy to fuel cellular processes. The study of mitochondria function and composition has implications for health and disease processes. Here, we present a new approach to comprehensively study the formation of large protein complexes of the electron transport system (the proteins responsible for energy production), which provides insight into mechanisms for increased mitochondrial efficiency with exercise training. This study deciphers the manner in which proteins assemble within complexes, and how this process is influenced by exercise training, a known physiological intervention to improve mitochondrial function and general health. Furthermore, we describe the regulation of additional protein complexes involved in mitochondrial function and health during exercise training.

## STAR★ Methods

### Key Resource Table

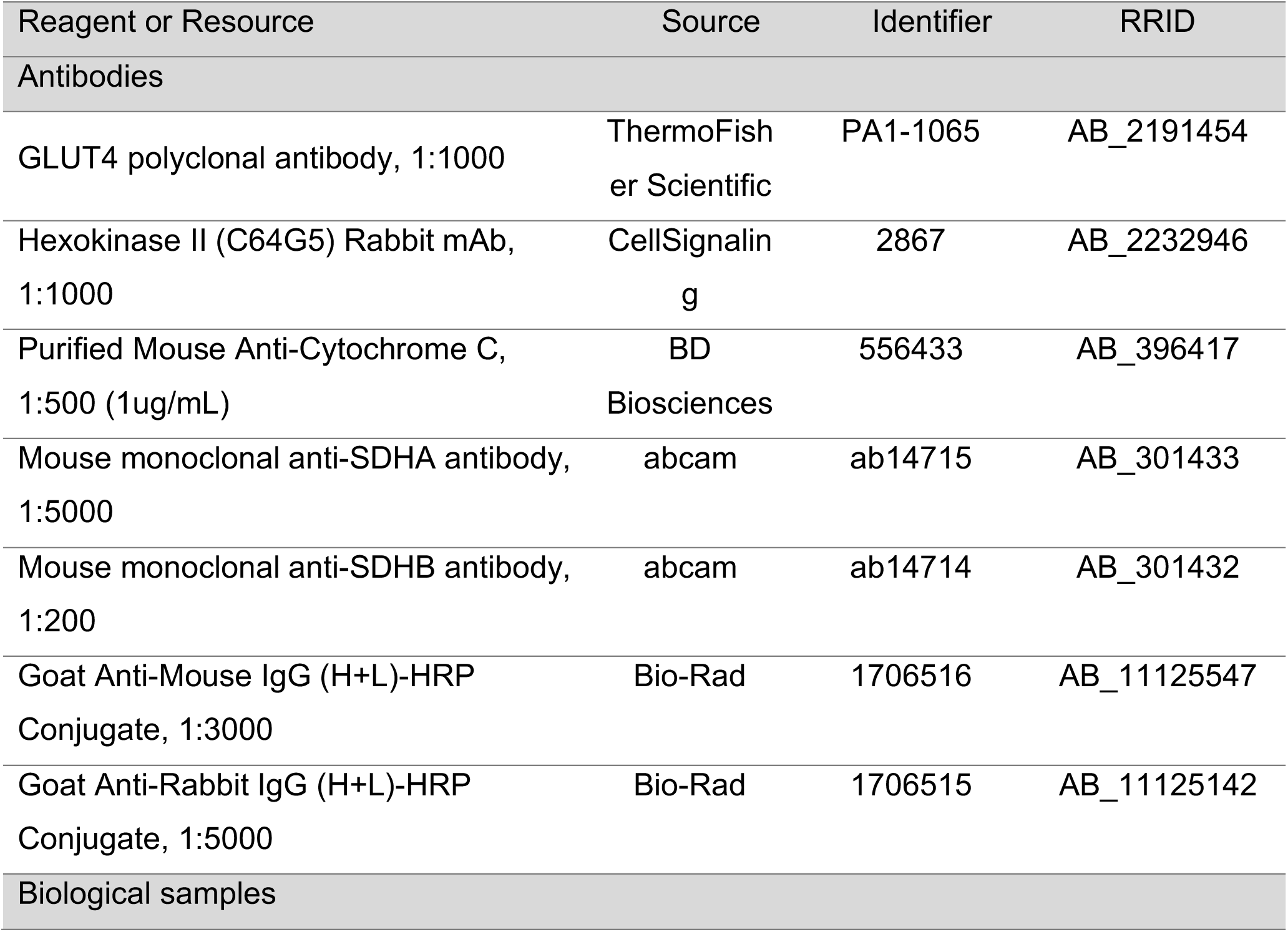
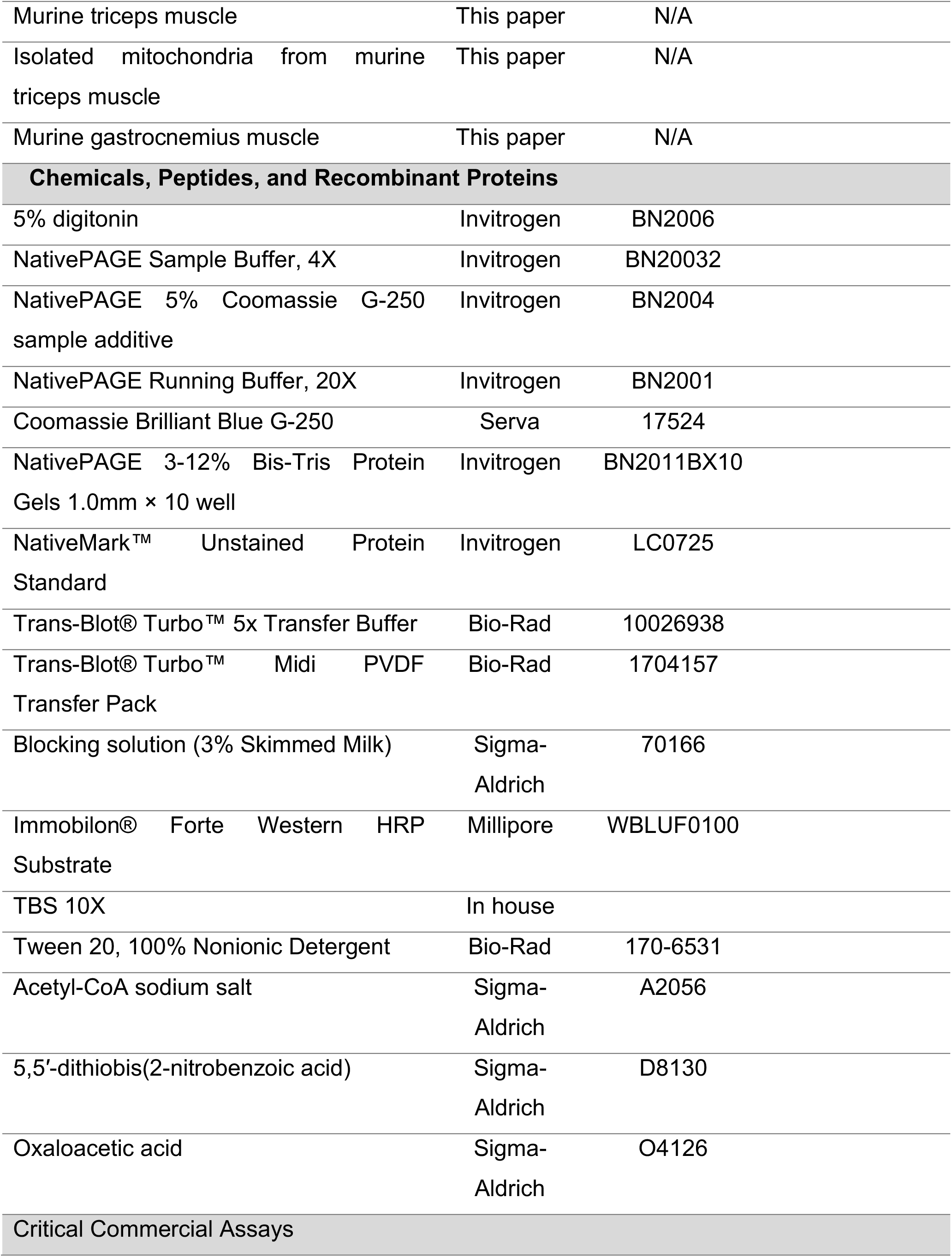
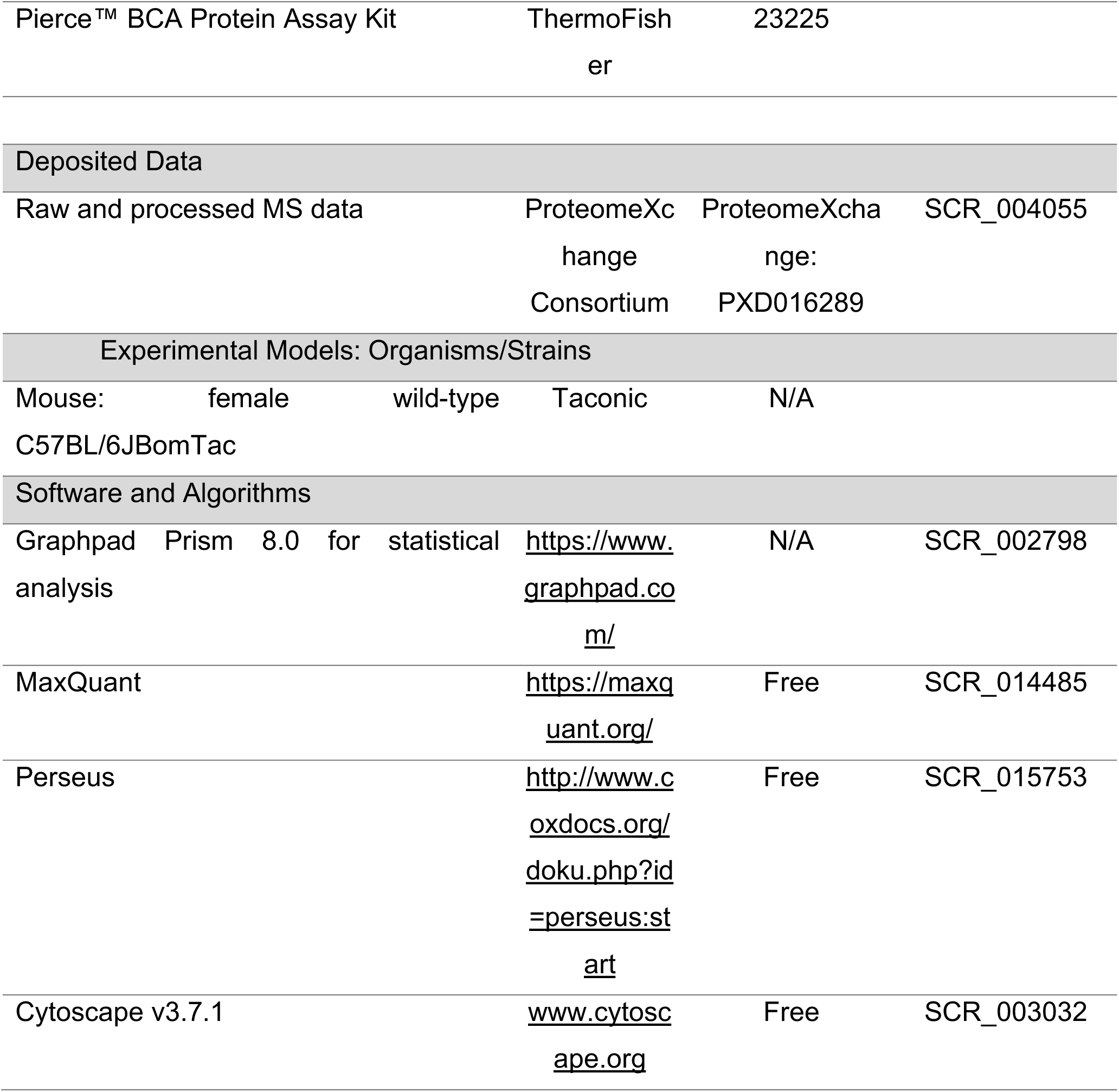

### Lead contact and materials availability

Requests for reagents and resources should be directed to the Lead Contact, Atul S. Deshmukh (Blegdamsvej 3B (07-8-19) - 2200 Copenhagen, Denmark).

### Animal and cell culture experiments

All animal experiments were approved by the Danish Animal Experimental Inspectorate and complied with the European Convention for the Protection of Vertebrate Animals Used for Scientific Purposes. Animals used in these experiments were 10- to 14-week old C57BL/6JBomTac female mice from Taconic and were kept on a 12:12 hour light-dark cycle with unlimited access to standard rodent diet and water *ad libitum*. Animals were divided into two experimental groups: sedentary and exercise trained. Mice were single-housed and the trained mice had access to a running wheel for 25 days. Running wheels were blocked 24h before they were euthanized at the end of the intervention, and *extensor digitorum longus*, *soleus*, *triceps brachii*, *gastrocnemius* and *quadriceps* skeletal muscle were collected. We observed larger exercise training effects in triceps muscle; as such this muscle group was selected for subsequent analyses. C2C12 muscle cells were grown and differentiated as described before (Deshmukh et al., 2015a).

### Sample preparation for total proteome of triceps muscle

Triceps muscles and differentiated C2C12 cells were lyzed in 0.1 M Tris-HCl, pH 7.5, 0.1 M DTT and 4% SDS, homogenized with an Ultra Turbax blender (IKA) and boiled for 5 min. The lysate was sonicated and clarified by centrifugation at 14000 rpm for 10 min. Samples were then processed following filter-aided sample preparation protocol using sequential endopeptidases LysC and trypsin for protein digestion (MED-FASP) (Wisniewski and Mann, 2012). Peptide fractions from LysC and trypsin digestion were then purified on C_18_ stage-tips (Rappsilber et al., 2003) prior to LCMS/MS analysis. The C2C12 muscle cells were included in the analysis to enhance protein identification by using the ‘match between runs’ algorithm in MaxQuant software (MaxQuant, RRID SCR_014485)(Schonke et al., 2018; Tyanova et al., 2016). The analysis was performed on n=5 biological replicates from each group (Sedentary, Training).

### Blue native polyacrylamide gel electrophoresis (BN-PAGE) and in gel digestion

Fresh triceps muscle was homogenized in mitochondria isolation buffer (100 mM sucrose, 100 mM KCl, 50 mM Tris-HCl, KH_2_PO_4_ 1 mM, 0.1 mM EGTA, 0.2% BSA) supplemented with the protease Nagarse. The lysate was cleared by low-speed centrifugation (750 g, 10 min), followed by high-speed centrifugation (10000 g, 10 min) to enrich for the mitochondrial fraction. The mitochondrial fraction was washed in isolation buffer and mitochondrial protein (50 μg) was prepared for electrophoresis on NativePAGE Novex 3-12% Bis-Tris Protein Gels (Invitrogen) as previously described (Jha et al., 2016). NativePAGE Sample Buffer (Invitrogen), 5% digitonin and 5% Coomassie G-250 was added to mitochondrial pellet and was electrophoresed at 150 V for 30 min at 4°C followed by 60 min at 250 V. Protein bands were visualized by Coomassie G-250 staining. Unstained marker bands (NativeMARK, Invitrogen) were visualized after fixation with 25% isopropanol and 10% acetic acid, and stained with 10% acetic acid and 60 mg/L of Coomassie R-250. Ten selected putative supercomplex bands, determined as described previously (Jha et al., 2016), were excised and cut into 1*1mm pieces followed by in-gel digestion as described (Shevchenko et al., 2006). Peptides from the trypsin digestion were then purified on C_18_ stage-tips (Rappsilber et al., 2007) before LCMS/MS analysis. The analysis was performed on n=4 biological replicates from each group (Sedentary, Exercise Training)

### LCMS/MS analysis

For the total proteome analysis, the peptides from the triceps muscles and C2C12s were measured with identical chromatographic conditions. Peptides were injected on a 50 cm C_18_-particle packed column (inner diameter 75 μm, 1.8 μm beads, Dr. Maisch GmbH, Germany) with Buffer A (0.5% formic acid) and separated over a 150-min linear gradient from 5-40% Buffer B (80% acetonitrile, 0.5% formic acid) at a flow rate of 250 nL/minute. The Easy nano-flow HPLC system was coupled to a LTQ Q-Exactive HF Orbitrap mass spectrometer via a nanoelectrospray source (all from Thermo Fisher Scientific, Germany). Mass spectra were acquired in a data-dependent manner, with automatic switching between MS and MS/MS using a top-12 method. MS spectra were acquired in the Orbitrap analyzer with a mass range of 300-1750 m/z and 60,000 resolutions at m/z 200. HCD peptide fragments acquired at 28 normalized collision energy were analyzed at high resolution in the Orbitrap analyzer. For the mitochondrial supercomplex proteome, peptides from in-gel digested samples were injected on a 15 cm C_18_-particle packed column (inner diameter 75 μm, 1.8 μm beads, Dr. Maisch GmbH, Germany) with Buffer A (0.5% formic acid) and separated over a 65-minute linear gradient from 5-40% Buffer B (80% acetonitrile, 0.5% formic acid) at a flow rate of 250 nL/minute. The Easy nano-flow HPLC system was coupled to a LTQ Q-Exactive HFX Orbitrap mass spectrometer via a nanoelectrospray source (all from Thermo Fisher Scientific, Germany). Mass spectra were acquired in a data-dependent manner, with automatic switching between MS and MS/MS using a top-12 method. MS spectra were acquired in the Orbitrap analyzer with a mass range of 300-1750 m/z and 60,000 resolutions at m/z 200. HCD peptide fragments acquired at 28 normalized collision energy were analyzed at high resolution in the Orbitrap analyzer.

### Mass spectrometry data analysis

Raw MS files were analyzed using MaxQuant software (https://www.maxquant.org/; RRID SCR_014485) (Tyanova et al., 2016). MS/MS spectra were searched by the Andromeda search engine (integrated into MaxQuant) against the decoy UniProt-mouse database supplemented with 262 frequently observed contaminants and forward and reverse sequences. In the main Andromeda search, precursor mass and fragment mass were matched with an initial mass tolerance of 6 and 20 ppm, respectively. The search included variable modification of methionine oxidation and N-terminal acetylation and fixed modification of carbamidomethyl cystein. Minimal peptide length was set to seven amino acids, and a maximum of two miscleavages were allowed. For the peptides and protein identification, the false discovery rate (FDR) was set to 0.01. MS runs from the triceps muscle and C2C12 myotubes were analyzed with the ‘match between runs’ option in MaxQuant. For matching, a retention time window of 30 s was selected. When all identified peptides were shared between two proteins, results were combined and reported as one protein group. In the case of myosin heavy chain proteins (Fig S1L), protein quantification was performed only with unique peptides.

### Protein quantification

For the triceps proteome, protein abundance (concentrations) were calculated based on the ‘Proteomics ruler’ concept (Wisniewski, 2017; Wisniewski et al., 2014). The raw protein intensities for individual proteins were divided by the summed signals of all proteins (on the basis of peptide identification) to obtain normalized total protein abundance. These values were further divided by the molecular weight of the proteins to yield the protein concentrations in pmol/mg of protein. Protein quantification for the mitochondrial supercomplexes was based on the MaxLFQ algorithm integrated into the MaxQuant software (RRID SCR_014485) (Cox et al., 2014).

### Bioinformatics analysis

Bioinformatics analysis was done in the Perseus software (http://www.coxdocs.org/doku.php?id=perseus:start; RRID SCR_015753). Categorical annotation was supplied in the form of Gene Ontology (GO) biological process (BP), molecular function (MF), and cellular component (CC). All annotations were extracted from the UniProt database. To retain sufficiently informative protein expression profiles for further analysis, the quantified proteins were filtered to have at least 3 valid values in at least one group (Sedentary, Training). The data was imputed to fill missing abundance values by drawing random numbers from a Gaussian distribution with a standard deviation of 30% and a downshift of 1.8 standard deviations from the mean. The imputed values have been tuned in order to simulate the distribution of low abundant proteins. The principal component analysis was performed on the imputed data matrix. For the total triceps proteome comparison between sedentary and the training groups, two samples *t*-test was performed (FDR = 0.05, s0 = 0.1). Results are presented as mean ± SEM. When comparing the mitochondrial proteome (BN-PAGE) between the sedentary and trained group, differentially expressed proteins were identified using an *a posteriori* information fusion scheme combining the biological relevance (fold-change) and the statistical significance (p-value) as described previously (Xiao et al., 2014). A ∏-value significance score cut-off of 0.05 was selected.

### Protein correlation profile (PCP)

The relative distribution of respiratory subunits across each band was calculated based on the PCP (Protein correlation profiling), a proteomics method for unbiased assignment of proteins to multiple fractions (often subcellular localizations) (Andersen et al., 2003; Foster et al., 2006; Krahmer et al., 2018). The protein abundance profiles from the analyzed gel bands was derived by scaling intensities for each quantified proteins over all bands within individual lane to a value of 0 to 1. Median PCP values for individual complexes within groups (sedentary and training) are also presented in Fig 3.

### Protein-protein interaction network analysis

Functional protein interaction networks were mapped using the STRING database (Szklarczyk et al., 2015) and further processed with Cytoscape (www.cytoscape.org; RRID SCR_003032). For the inclusion of an interaction in the network mapping a very high confidence score (0.7) was required. For the network analysis, all oxidative phosphorylation proteins, electron carriers and mitochondrial respiratory complex assembly factors were included for the bands analyzed.

### Immunoblot analysis

Western blot was performed with lyzed samples from muscles *extensor digitorum longus*, *soleus*, *triceps*, *gastrocnemius* and *quadriceps* for immunoblotting for GLUT4 (#PA1-1065 Thermo Fisher, RRID AB_2191454), hexokinase II (HKII, Cell Signalling, 2867, RRID AB_2232946) and CytC (BD Biosciences #556433, RRID AB_396417).

For BN-PAGE, after electrophoresis, the complexes were transferred onto PVDF membranes (semidry transfer, BioRad Trans-Blot Turbo, high molecular weight program, 10 min) and probed with specific antibodies against Sdha (abcam ab14715, RRID AB_301433) and Sdhb (abcam ab14714, RRID AB_301432). Chemiluminescent membranes were imaged using the ChemiDoc XRS+ (Bio-Rad, CA, USA).

### Citrate synthase activity assay

Gastrocnemius muscle was pulverized, lyzed and centrifuged prior to supernatant collection. The citrate synthase activity assay (based on (Alp et al., 1976)) consisted on a citrate synthase activity reaction mix (0.1 M This HCl, pH 8.1, 0.4 mM acetyl-CoA sodium salt (Sigma-Aldrich) and 1 μM 5,5′-dithiobis(2-nitrobenzoic acid) (Sigma-Aldrich)) added to muscle protein extract prior to absorbance measurement (412 nm) every 30 s for 5 min (basal slope) followed by addition of oxaloacetic acid (10 mM) and remeasurements every 30 s for another 5 min (reaction slope). The citrate synthase activity was calculated from the delta of the reaction and basal slopes and is presented in μmol/min/g protein.

### Data availability

The mass spectrometry proteomics data have been deposited to the ProteomeXchange Consortium (SCR_004055) via the PRIDE partner repository (Perez-Riverol et al., 2019) with the dataset identifier PXD016289.

**Figure S1.**
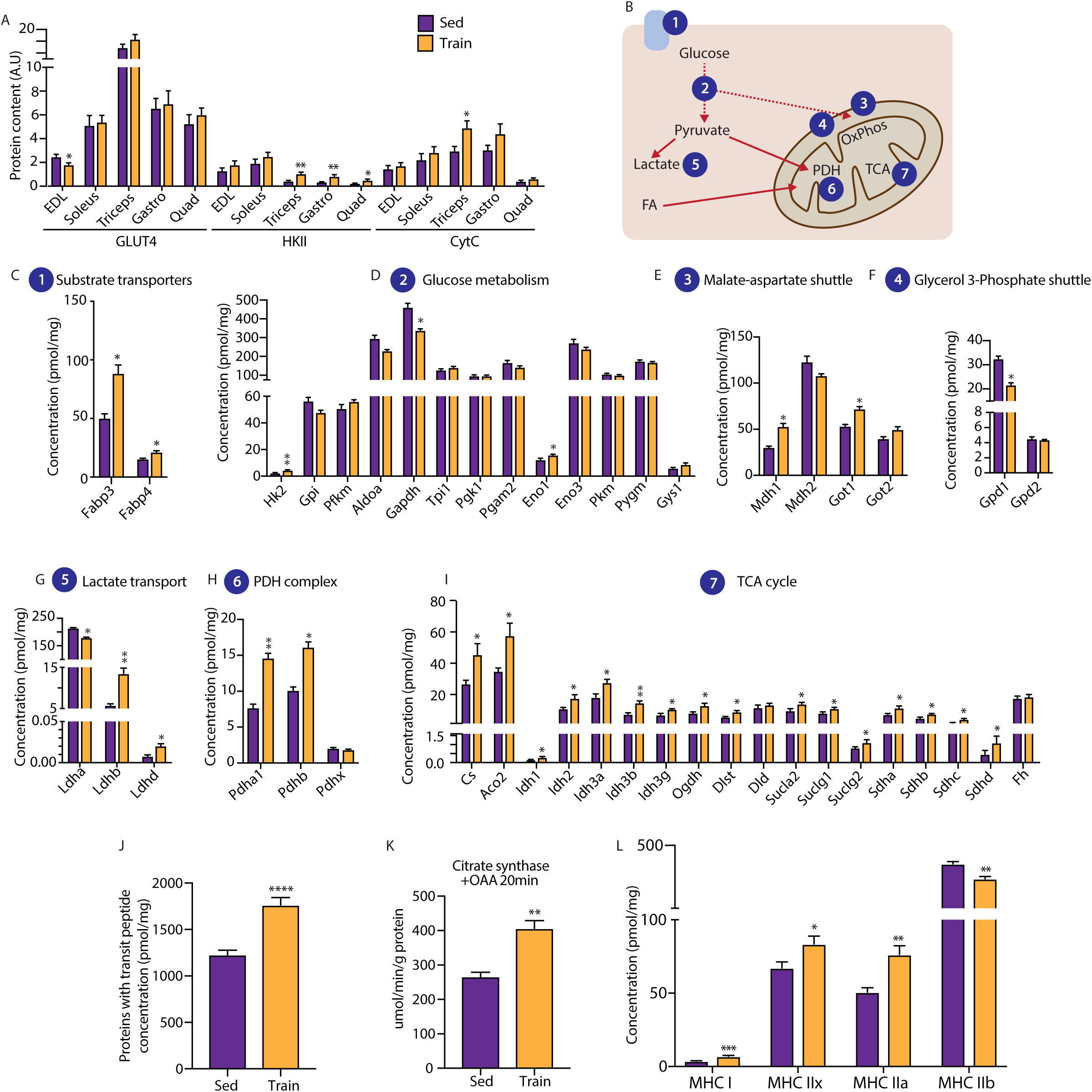
A: Western blotting-based quantification of GLUT4, HKII and CytC protein levels in sedentary and trained mice (n=5 per group). B: Schematic of different pathways studied (C-I). C-I: Protein concentration in the triceps proteome for substrate transporter proteins (C), glucose metabolism proteins (D), malate-aspartate shuttle proteins (E), glycerol-3-phosphate shuttle proteins, (F), lactate transport proteins (G), pyruvate dehydrogenase (PDH) complex proteins (H), and tricarboxylic acid (TCA) cycle proteins (I). J-K: Total protein concentration in the triceps proteome for proteins with transit peptides (J). K: Citrate synthase activity by addition of oxaloacetate (OAA). L: Protein concentration in the triceps proteome for fiber type specific proteins. All data is represented as mean ± SEM. *p<0.05, **p<0.01, ***p<0.001, ****p<0.0001. Sed: sedentary; Train: exercise trained; HK: hexokinase 2.

TableS1. Proteins quantified in triceps muscle proteome (n=5 per group).

https://drive.google.com/open?id=1dTyasnzEpu2G5v0LqI-gjBTB2Ac3zeAj

TableS2. Mitochondrial proteins quantified from BN-PAGE gel bands (n=4 per group).

https://drive.google.com/open?id=1dTyasnzEpu2G5v0LqI-gjBTB2Ac3zeAj

TableS3. Protein correlation profile values for oxidative phosphorylation proteins

https://drive.google.com/open?id=11DWzg9f5K3MkGpcF3u1-Lu9-kEKXIQ-S

TableS4. Protein-protein interaction matrix from oxidative phosphorylation proteins in band 7.

https://drive.google.com/open?id=1_WggKgr6z1A-lrTcmJG56KOy8F-VVPAh

TableS5. Protein-protein interaction matrix from oxidative phosphorylation proteins in band 2.

https://drive.google.com/open?id=1d3o6FseD0MBgBuuJL4bw0ZvdSJCi-Ezu

